# From smartphones to satellites: Uniting crowdsourced biodiversity monitoring and Earth observation to fill the gaps in global plant trait mapping

**DOI:** 10.1101/2025.03.10.641660

**Authors:** Daniel Lusk, Sophie Wolf, Daria Svidzinska, Carsten F. Dormann, Jens Kattge, Helge Bruelheide, Francesco Maria Sabatini, Gabriella Damasceno, Álvaro Moreno Martínez, Cyrille Violle, Daniel Hending, Georg J. A. Hähn, Solana Tabeni, Shyam Phartyal, Fernando Gonçalves, Holger Kreft, Marco Schmidt, Han Chen, Behlül Güler, Jiri Dolezal, Remigiusz Pielech, Anaclara Guido, Ciara Dwyer, Francesca Napoleone, Jacob Willie, André Luís Gasper, Manuel J. Macía, Milan Chytry, Jonathan Lenoir, Dinesh Thakur, Jürgen Dengler, Sebastian Świerszcz, Jan Altman, Ladislav Mucina, Ashish N. Nerlekar, Kaoru Kakinuma, Pravin Rawat, Zvjezdana Stančić, Riccardo Testolin, Mohamed Z. Hatim, Flávio Rodrigues, Jürgen Homeier, Marcia C. M. Marques, James K. McCarthy, M. A. El-Sheikh, Teja Kattenborn

## Abstract

Plant functional traits are fundamental to ecosystem dynamics and Earth system processes, but their global characterization is limited by the availability of field surveys and trait measurements. Recent expansions in biodiversity data aggregation, including large collections of vegetation surveys, citizen science observations, and trait measurements, offer new opportunities to overcome these constraints. Here we demonstrate that combining these diverse data sources with high-resolution Earth observation data enables accurate modeling of key plant traits at up to 1 km resolution. Our approach achieves high predictive power, reaching correlations up to 0.63 (15 of 31 traits exceeding 0.50) and improved spatial transferability, effectively bridging gaps in under-sampled regions. By capturing a broad range of traits with high spatial coverage, these maps can enhance our understanding of plant community properties and ecosystem functioning globally, and can serve as useful tools in modeling global biogeochemical processes and informing worldwide conservation efforts. Ultimately, our framework highlights the power and necessity of crowdsourced biodiversity data in high-resolution plant trait modeling. We anticipate that advancements in biodiversity data collection and remote sensing capabilities will further refine global trait mapping, fostering a dynamic trait-based understanding of the biosphere.

## 1 Introduction

The diversity and distribution of plants shape ecosystem function and influence global biogeochemical cycles [1–3]. Since Humboldt’s early explorations of plant geography in 1807, ecologists have sought to understand how plant properties govern species’ success across environmental gradients and their role in Earth system processes [4–7]. Despite significant advances, a comprehensive global understanding of plant functional traits and their implications for ecosystem resilience remains incomplete [8, 9].

Plant functional traits—measurable characteristics influencing growth, reproduction, and survival—are central to ecological strategies and ecosystem modeling [10–13]. Traits such as leaf area, wood density, and seed mass reflect trade-offs in resource use, stress tolerance, and competition, shaping plant community assembly [14]. Integrating these traits into global vegetation and Earth system models is crucial for refining projections of energy, carbon, and water cycles [15, 16]. However, producing spatially explicit, high-resolution trait maps requires extensive in situ trait measurements—a challenge that remains largely unmet.

Earth observation technologies provide valuable insights into vegetation properties, offering continuous, high-resolution data on surface reflectance, vegetation structure, climate, and soil conditions [17–22]. Yet, their utility for plant trait modeling remains constrained by limited in situ trait observations for calibration and validation. While in situ trait measurements are accurate, they are geographically sparse and labor-intensive [23]. Global initiatives such as TRY and BIEN have compiled extensive trait databases [24–26], but they often lack the spatial coverage necessary for coherently mapping plant functional composition at fine scales. Consequently, satellite-based trait extrapolations carry significant uncertainties [27, 28].

Crowdsourced ecological data offer a promising avenue to address these gaps. Expert-led vegetation surveys, such as those aggregated by the sPlot database, document species co-occurrence and abundance around our planet [29–31]. However, many regions remain underrepresented (Fig. 2. Meanwhile, citizen science platforms like iNaturalist and Pl@ntNet, aggregated through biodiversity databases like the Global Biodiversity Information Facility (GBIF), have contributed over half a billion plant observations (Fig. 2), vastly expanding global species distribution datasets [32–34]. Though these data exhibit spatial and taxonomic biases, they hold great potential for plant trait inference. Recent work by Wolf *et al.* [28] demonstrated that linking species occurrences from citizen science platforms with trait databases can approximate community-weighted mean trait distributions with remarkable accuracy.

Each data source—trait measurements, vegetation surveys, and citizen science observations— has strengths and limitations. Here, we present a novel data integration framework comparing and combining these sources with high-resolution Earth observation datasets to generate continuous global maps of 31 key plant traits. Our approach integrates environmental data on surface reflectance, climate, soil properties, and vegetation structure with species observations from GBIF, expert vegetation surveys from sPlot, and trait records from TRY to improve spatial coverage, resolution, and accuracy (Fig. 1).

**Figure 1:**
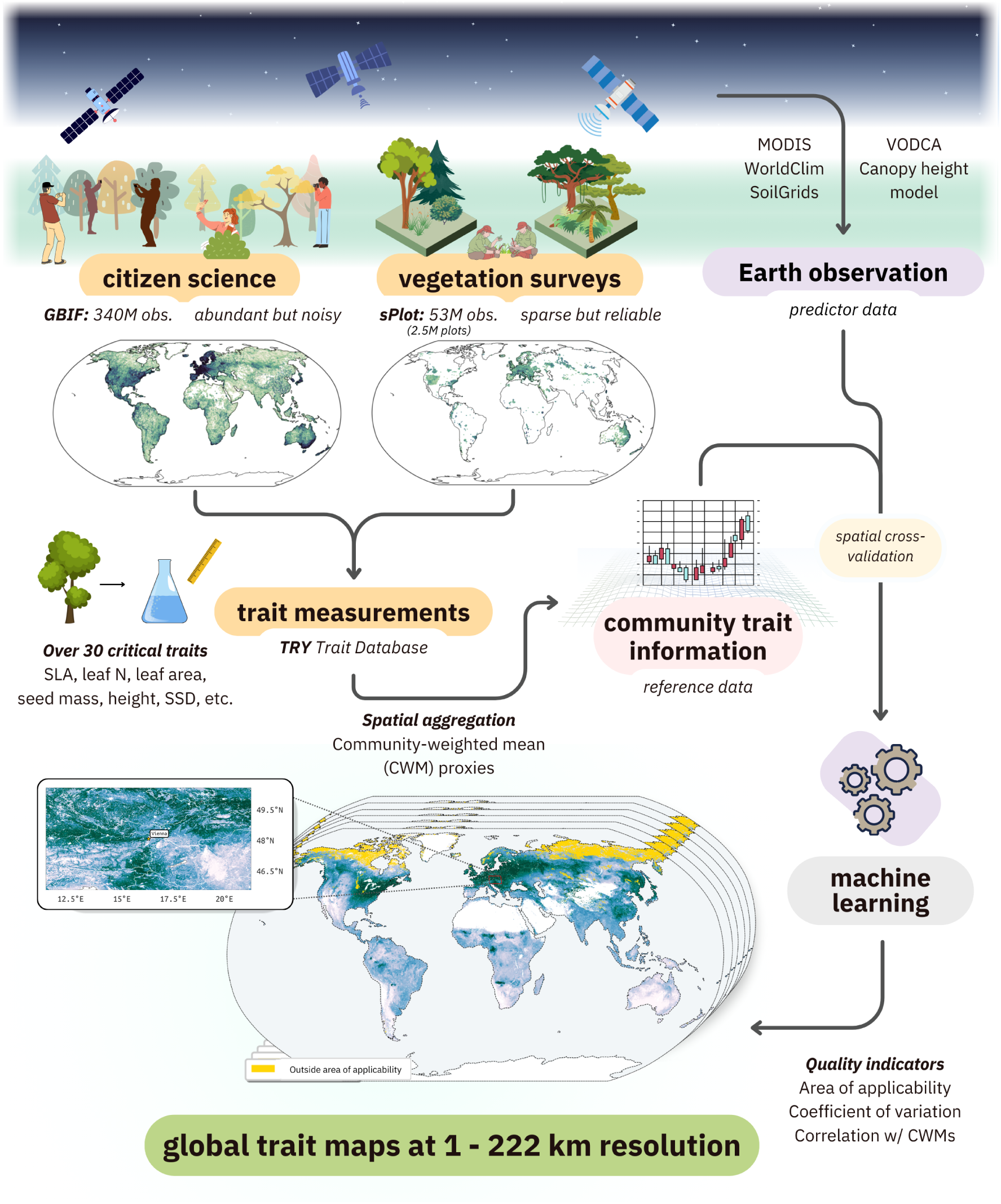
Workflow for modeling plant functional traits using a combination of diverse data sources, including citizen science observations, expert-organized vegetation surveys, curated trait measurement data, and Earth observation-derived predictor variables. The approach integrates community composition and trait information as reference data to train machine learning models, which generate global community-weighted mean trait maps at up to 1 km resolution.

We predict community-weighted mean trait values globally using three data subsets: (1) scientific vegetation surveys alone (SCI), (2) citizen science observations alone (CIT), and (3) a combined approach (COMB). Using spatial cross-validation with independent vegetation survey data, we assess model performance and geographic generalizability across scales from 1 km to 222 km. Our approach achieves unprecedented accuracy for many traits and, to our knowledge, maps several traits at high resolution for the first time. By integrating diverse data sources, we produce the most precise large-scale trait maps to date, significantly improving spatial coverage and predictive reliability. These advances provide a robust foundation for refining ecosystem models and predicting global vegetation dynamics with greater confidence.

## 2 Results

### 2.1 Density and spatial coverage of citizen science and vegetation survey data

The three trait data subsets—vegetation surveys (SCI), citizen science observations (CIT), and their combination (COMB)—exhibited distinct patterns of observation density and geographic coverage. Prior to matching with trait measurements from the TRY trait database, CIT contained 339,971,350 observations of 314,217 species, while SCI consisted of 2,534,183 plots recording 52,942,365 observations of 91,603 species. After matching with TRY, CIT retained 89% of observations (*n* = 303,288,097) and 28% of species (*n* = 88,802), while SCI retained consisting of 84% of plot-level relative abundance (*n* = 45,037,375 observations) and 43% of species (*n* = 39,286). Following spatial aggregation for matching with environmental predictors, CIT (and therefore COMB) displayed markedly greater spatial coverage than SCI alone, though both subsets showed spatial clustering in Europe, North America, Japan, and Australia (Fig. 2).

**Figure 2:**
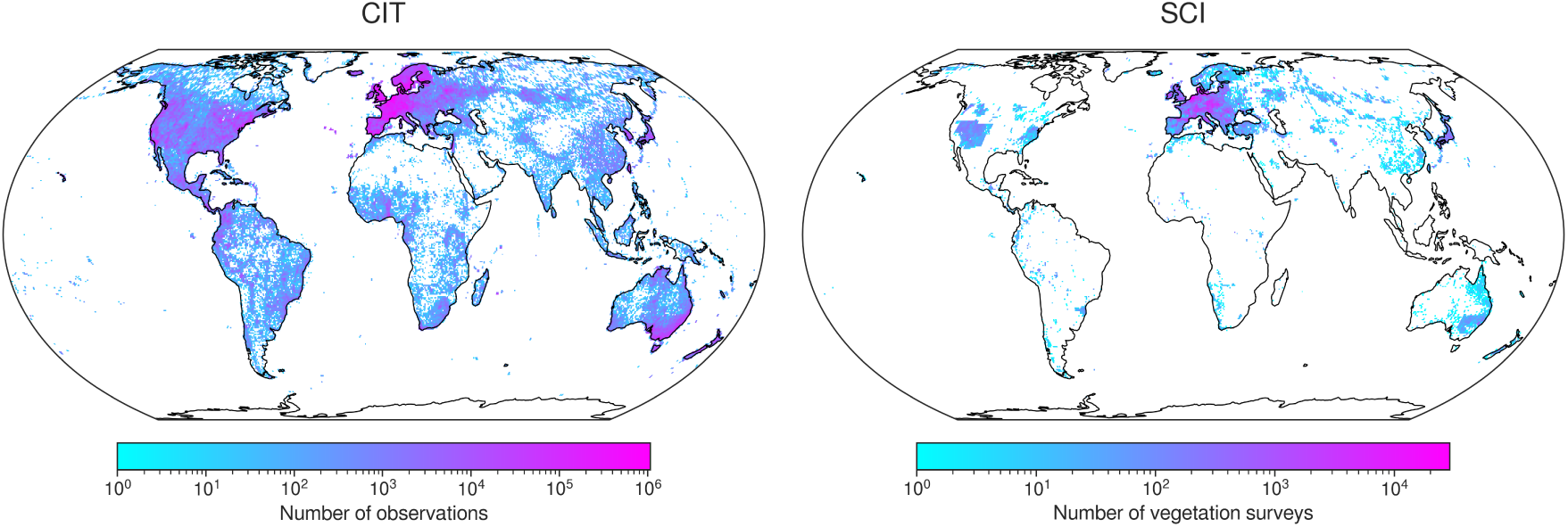
Density maps of citizen science observations (CIT) and vegetation surveys (SCI) after matching with trait data from the TRY trait database, spatial aggregation, and filtering.

### 2.2 Global trait maps and spatial transferability

We modeled the global distributions of 31 plant traits (Table 2) across five spatial resolutions: 1 km (∼0.01°), 22 km (∼0.2°), 55 km (∼0.5°), 111 km (∼1°), and 222 km (∼2°) (Fig. 3). To account for differences in observation density and geographic coverage, models were generated for each of the three trait data subsets: SCI, CIT, and COMB. We used gradient-boosted decision trees, which are well suited for capturing complex, non-linear relationships, with environmental predictors aggregated at each corresponding spatial resolution [35]. Rather than deriving coarser-resolution maps from higher-resolution outputs, we trained separate models at each scale to prevent biases from simple upscaling. This approach yielded a total of 465 models (31 traits × 5 resolutions × 3 trait data subsets).

**Figure 3:**
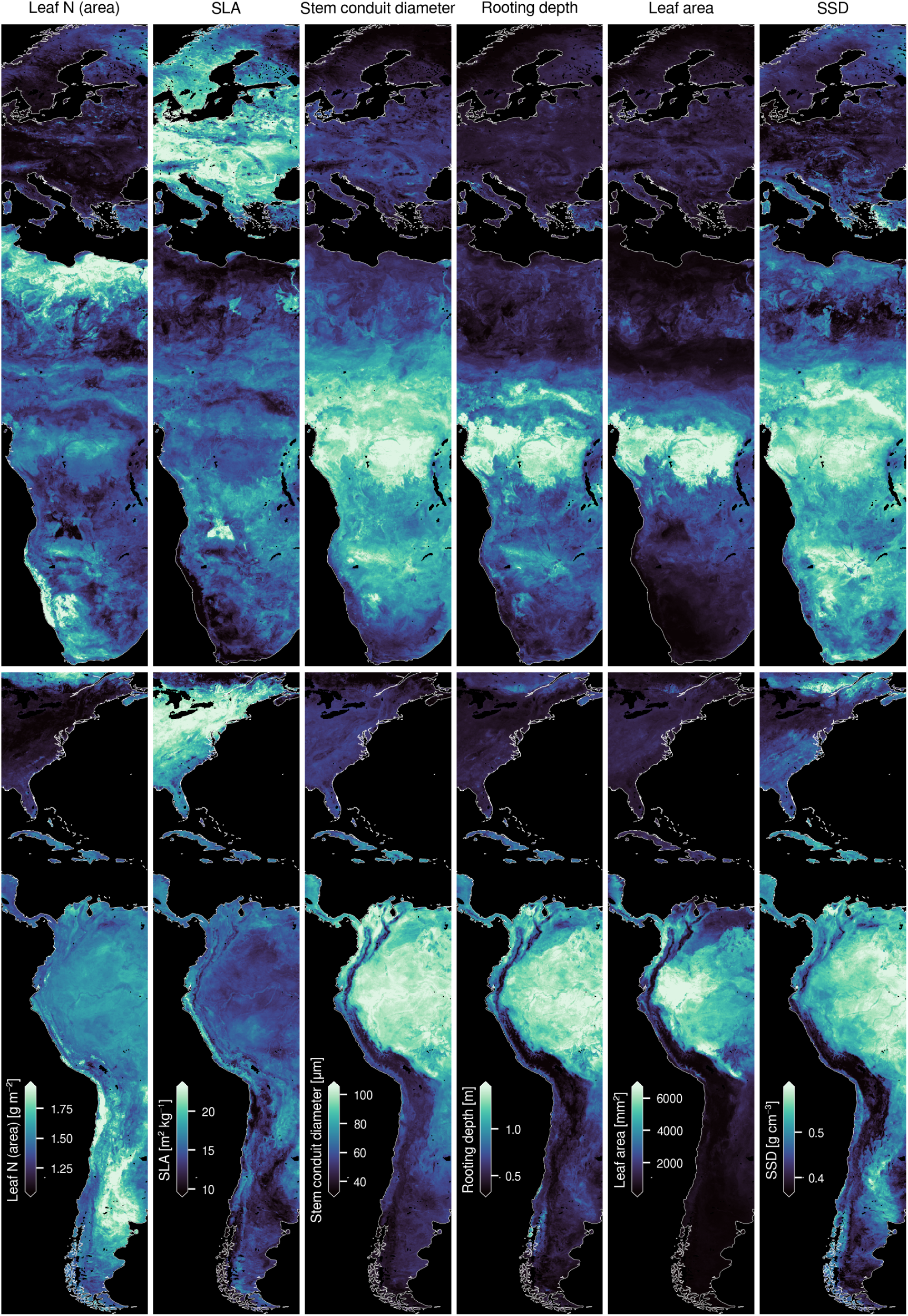
Trait maps based on combined citizen science and vegetation survey data (COMB) at 1 km resolution across two longitudinal sections of Earth: northern Europe to southern Africa (top row) and eastern North America to western South America (bottom row). Maps are presented here in WGS84 projection (EPSG:4326), and full, equal area trait maps can be **viewed online** or downloaded directly (see Data Availability).

**Table 1:** Pearson’s correlation coefficient *r* of each trait map in relation to spatially-independent gridded sPlot community-weighted means (CWM) unused in model training at different resolutions (1 km - 222 km). The highest correlation for each trait-resolution pair is in bold and the second-highest is in italics. Models incorporating a combination of citizen science and vegetation survey data (COMB) had the strongest correspondence for all traits and all resolutions, and models incorporating citizen science data only (CIT) exhibited the second-strongest correlations for most trait-resolution pairs. All trait values were transformed using a Yeo-Johnson power transform with the same parameters determined in the original transformation of the sPlot CWMs.

| Author | Resolution | SLA | Leaf N (mass) | Leaf N (area) |
| --- | --- | --- | --- | --- |
| This study (COMB) | 1 km | **0.63** | **0.56** | **0.63** |
|  | 22 km | **0.65** | **0.62** | **0.68** |
|  | 55 km | **0.63** | **0.59** | **0.65** |
|  | 111 km | **0.59** | **0.54** | **0.60** |
|  | 222 km | **0.52** | **0.46** | **0.59** |
| This study (CIT) | 1 km | *0.53* | *0.49* | *0.55* |
|  | 22 km | *0.45* | *0.49* | *0.53* |
|  | 55 km | *0.44* | *0.47* | *0.54* |
|  | 111 km | *0.42* | *0.44* | *0.52* |
|  | 222 km | *0.41* | *0.37* | *0.51* |
| van Bodegom et al., 2014 | 55 km | 0.33 | - | - |
|  | 111 km | 0.23 | - | - |
|  | 222 km | 0.24 | - | - |
| Boonman et al., 2020 | 55 km | 0.43 | 0.11 | 0.44 |
|  | 111 km | 0.37 | 0.08 | 0.42 |
|  | 222 km | 0.37 | 0.12 | 0.41 |
| Butler et al., 2017 | 55 km | 0.29 | 0.20 | 0.39 |
|  | 111 km | 0.36 | 0.29 | 0.37 |
|  | 222 km | 0.29 | 0.25 | 0.33 |
| Madani et al., 2018 | 55 km | 0.10 | - | - |
|  | 111 km | 0.25 | - | - |
|  | 222 km | 0.25 | - | - |
| Moreno et al., 2018 | 1 km | 0.38 | 0.26 | - |
|  | 22 km | 0.38 | 0.31 | - |
|  | 55 km | 0.38 | 0.12 | - |
|  | 111 km | 0.40 | 0.21 | - |
|  | 222 km | *0.44* | 0.21 | - |
| Schiller et al., 2021 | 55 km | *0.47* | 0.38 | 0.50 |
|  | 111 km | 0.38 | 0.35 | 0.46 |
|  | 222 km | 0.39 | 0.31 | 0.49 |
| Vallicrosa et al., 2022 | 1 km | - | 0.29 | - |
|  | 22 km | - | 0.37 | - |
|  | 55 km | - | 0.20 | - |
|  | 111 km | - | 0.33 | - |
|  | 222 km | - | 0.30 | - |
| Wolf et al., 2022 | 22 km | 0.41 | 0.31 | 0.38 |
|  | 55 km | 0.42 | 0.27 | 0.41 |
|  | 111 km | 0.32 | 0.29 | 0.34 |
|  | 222 km | 0.31 | 0.36 | 0.37 |

**Table 2:**
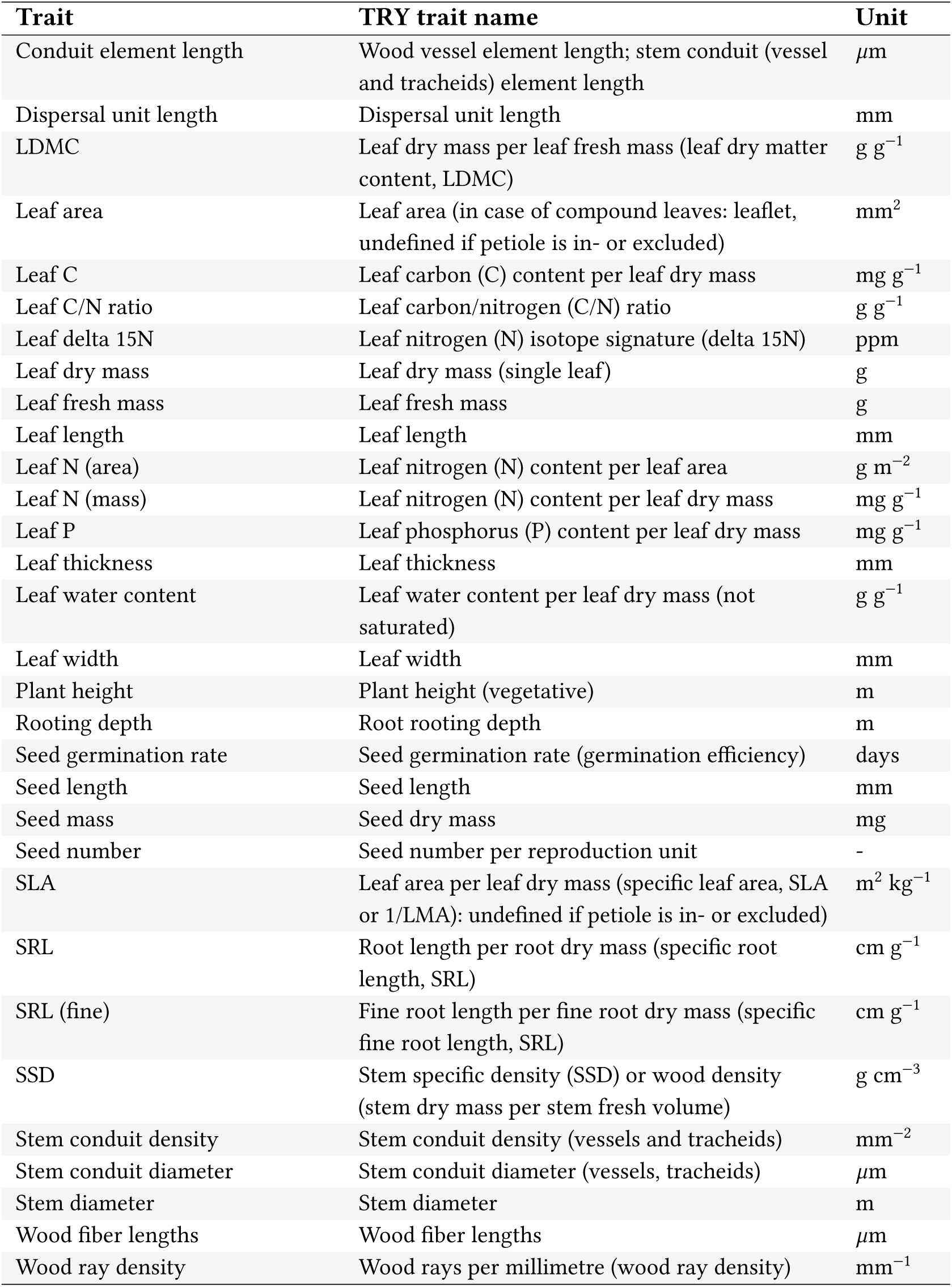
Plant functional traits used for modeling and trait map generation. “Trait” refers to a shortened version of the “TRY trait name”. “TRY trait name” refers to the name used in the TRY trait database.

| **Trait** | **TRY trait name** | **Unit** |
| --- | --- | --- |
| Conduit element length | Wood vessel element length; stem conduit (vessel and tracheids) element length | $\mu\text{m}$ |
| Dispersal unit length | Dispersal unit length | mm |
| LDMC | Leaf dry mass per leaf fresh mass (leaf dry matter content, LDMC) | $\text{g g}^{-1}$ |
| Leaf area | Leaf area (in case of compound leaves: leaflet, undefined if petiole is in- or excluded) | $\text{mm}^2$ |
| Leaf C | Leaf carbon (C) content per leaf dry mass | $\text{mg g}^{-1}$ |
| Leaf C/N ratio | Leaf carbon/nitrogen (C/N) ratio | $\text{g g}^{-1}$ |
| Leaf delta 15N | Leaf nitrogen (N) isotope signature (delta 15N) | ppm |
| Leaf dry mass | Leaf dry mass (single leaf) | g |
| Leaf fresh mass | Leaf fresh mass | g |
| Leaf length | Leaf length | mm |
| Leaf N (area) | Leaf nitrogen (N) content per leaf area | $\text{g m}^{-2}$ |
| Leaf N (mass) | Leaf nitrogen (N) content per leaf dry mass | $\text{mg g}^{-1}$ |
| Leaf P | Leaf phosphorus (P) content per leaf dry mass | $\text{mg g}^{-1}$ |
| Leaf thickness | Leaf thickness | mm |
| Leaf water content | Leaf water content per leaf dry mass (not saturated) | $\text{g g}^{-1}$ |
| Leaf width | Leaf width | mm |
| Plant height | Plant height (vegetative) | m |
| Rooting depth | Root rooting depth | m |
| Seed germination rate | Seed germination rate (germination efficiency) | days |
| Seed length | Seed length | mm |
| Seed mass | Seed dry mass | mg |
| Seed number | Seed number per reproduction unit | - |
| SLA | Leaf area per leaf dry mass (specific leaf area, SLA or 1/LMA): undefined if petiole is in- or excluded) | $\text{m}^2 \text{kg}^{-1}$ |
| SRL | Root length per root dry mass (specific root length, SRL) | $\text{cm g}^{-1}$ |
| SRL (fine) | Fine root length per fine root dry mass (specific fine root length, SRL) | $\text{cm g}^{-1}$ |
| SSD | Stem specific density (SSD) or wood density (stem dry mass per stem fresh volume) | $\text{g cm}^{-3}$ |
| Stem conduit density | Stem conduit density (vessels and tracheids) | $\text{mm}^{-2}$ |
| Stem conduit diameter | Stem conduit diameter (vessels, tracheids) | $\mu\text{m}$ |
| Stem diameter | Stem diameter | m |
| Wood fiber lengths | Wood fiber lengths | $\mu\text{m}$ |
| Wood ray density | Wood rays per millimetre (wood ray density) | $\text{mm}^{-1}$ |

Final trait maps, along with each map’s corresponding coefficient of variation (COV) and area of applicability (AOA) masks, were generated as GeoTIFF rasters. COMB maps at 1 km resolution can be viewed online via an **interactive map viewer** or downloaded directly (see Data Availability). Predictions were made for all pixels where sufficient predictor data were present with the exclusion of permanent water bodies, and map users are encouraged to consider their specific use cases alongside the provided model performance and uncertainty indicators (COV and AOA). Plant functional type-specific maps are also available for download.

#### 2.2.1 Effects of incorporating citizen science data on spatial transferability

Citizen science observations, with their significantly greater geographic extent and sample size, have the potential to enhance vegetation survey data by allowing models to predict with greater confidence in data-deficient regions. To evaluate this, we calculated two metrics for each trait map and model during spatial cross-validation: area of applicability [36] and coefficient of variation (see Methods) (Fig. 4). For this portion of the analysis, 1 km maps were primarily considered to ensure comparison at the finest grain size available, though area of applicability was also compared across resolutions.

**Figure 4:**
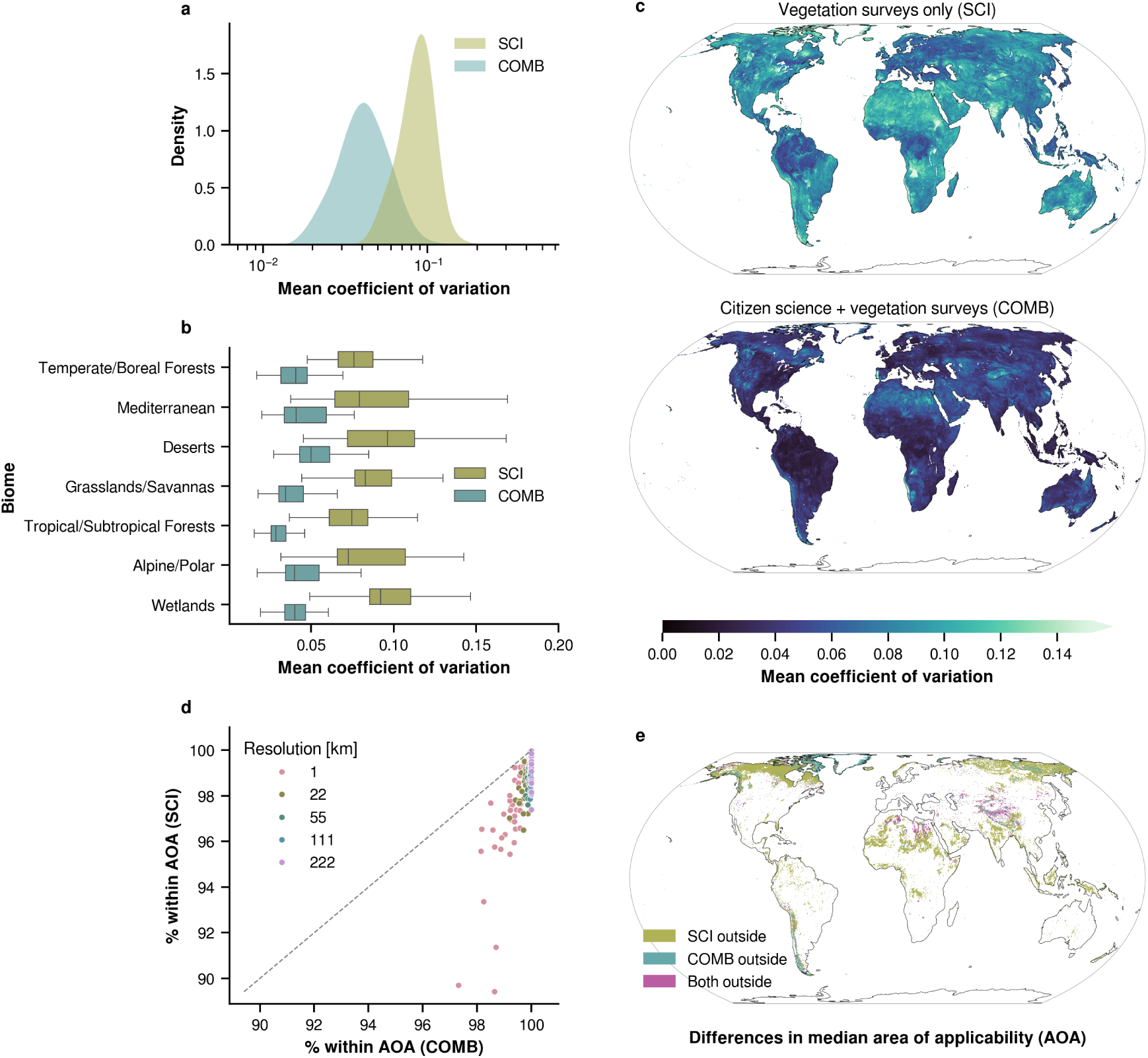
Spatial transferability metrics for traits with *r* ≥ 0.5 at 1 km resolution. **a.** Mean coefficient of variation (COV) from spatial cross-validation for models using only vegetation survey trait data (SCI) and those combining vegetation survey and citizen science data (COMB). **b.** Mean COV for SCI and COMB models by biome. Boxes indicate the 25th to 75th percentiles, with whiskers showing the interquartile range. See Table A.2 for more information on biome definition. **c.** Pixel-wise mean COV for SCI (top) and COMB (bottom). Including citizen science data generally lowered the pixel-wise COV compared to using vegetation survey data alone. **d.** Scatter plot of the area of applicability (AOA) for SCI vs. COMB models across different resolutions. The percentage of prediction pixels within the AOA is shown, with colors representing model resolution. A one-to-one line is included for reference. COMB models consistently showed a larger AOA than SCI across all resolutions. **e.** Pixel-wise median AOA for SCI and COMB. Olive areas indicate where only SCI predictions are unreliable, blue where only COMB predictions are unreliable, and magenta where both are outside the AOA.

The area of applicability (AOA) of a model refers to its ability to be applied to new regions, determined by how similar the new region’s predictors are to those used during the model’s training [36]. Incorporating citizen science data alongside vegetation surveys increased the AOA for all traits at all resolutions, with an average gain of 2.43 percentage points. The largest improvement was 9.22 for wood fiber length at 1 km resolution, while the smallest was 0.04 for leaf width at 222 km resolution. In particular, trait maps at finer resolutions benefited more on average from the inclusion of citizen science trait data, while the benefits became smaller at coarser resolutions (Fig. 4d).

The coefficient of variation (COV) is another indicator of model uncertainty derived by comparing the predictions made by each model during cross-validation. For all models, the COV was consistently lower on average for COMB maps than for SCI maps. In particular, COMB maps showed better generalization across all biomes, including regions where observations are especially sparse for both citizen science observations and vegetation surveys such as desert and alpine areas.

### 2.3 Model performance across trait data subsets at 1 km resolution

To assess predictive accuracy at high spatial resolution, we compared model performance across the three trait data subsets (SCI, CIT, and COMB) at 1 km resolution, using spatial cross-validation with independent vegetation survey community-weighted mean (CWM) trait data not employed in model training (Fig. 5a-c).

**Figure 5:**
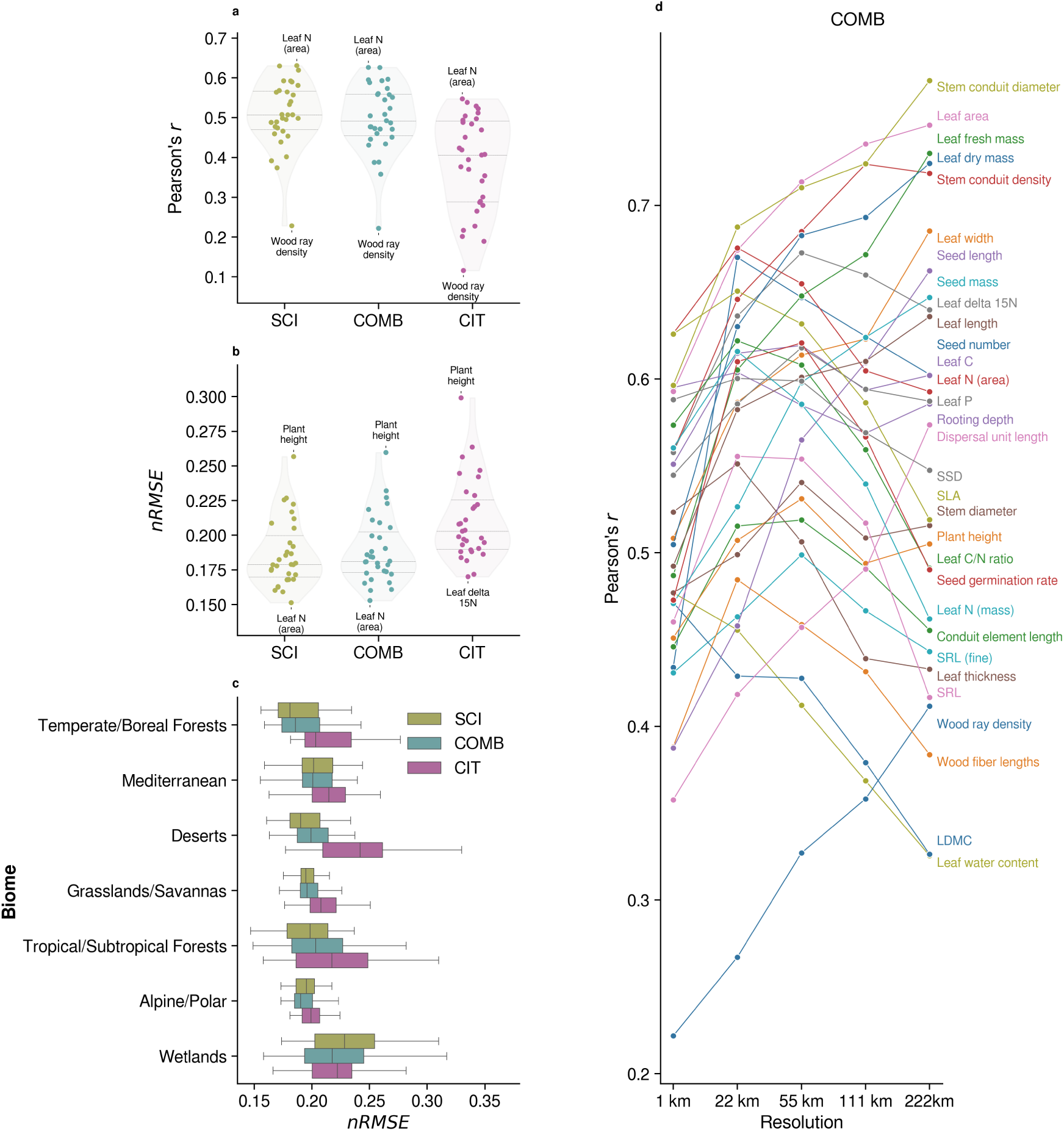
Model performance across trait data subsets: vegetation survey data alone (SCI), citizen science data alone (CIT), and a combination of the two (COMB). All metrics are the result of validation against spatially independent, held-out vegetation survey community-weighted mean trait values. **a., b.** Pearson correlation coefficient *r* (a) and normalized root mean squared error *nRMSE* (b) of trait models for SCI, COMB, and CIT trait data subsets. The points indicate individual trait model performance, while the encompassing violin plots represent the performance distribution for each trait subset. Inner horizontal bars represent the 75th, 50th (median), and 25th quantiles. **c.** *nRMSE* of SCI, COMB, and CIT models across biomes. See Table A.2 for more information about biome selection. **d.** Pearson’s correlation coefficient *r* for COMB models trained on data aggregated at five spatial resolutions.

SCI and COMB models consistently outperformed CIT models in terms of correlation coefficient (*r*) and normalized root mean squared error (nRMSE). COMB models maintained similar *r* and nRMSE values to those trained solely on SCI data, with an average increase in *r* by 0.12 compared to CIT models. At 1 km resolution, SCI models had *r* ≥ 0.5 for 16 out of 31 traits, COMB models for 15 traits, and CIT models for 6 traits. The traits with the highest predictive accuracy across both COMB and SCI models included leaf N (area), specific leaf area (SLA), rooting depth, stem specific density (SSD), stem conduit diameter, and leaf area. Notably, COMB models displayed robust predictive accuracy across biomes, performing similarly to SCI models even in data-sparse regions.

### 2.4 Map accuracy of combined trait data at different spatial scales

To evaluate how spatial resolution affected map accuracy, we compared Pearson’s *r* for COMB models across five resolutions (1 km - 222 km), where training data was aggregated at each resolution prior to model training to avoid simple upscaling effects (Fig. 5d). Accuracy generally improved with increasing spatial aggregation, with most traits (18 of 31) showing peak performance at 22 km or 55 km resolution. Beyond this point, accuracy tended to plateau or slightly decline, suggesting diminishing predictability at coarser scales for many traits. Similar trends were observed for SCI and CIT models (see Appendix A.4).

### 2.5 Comparison with existing trait map products

To benchmark model performance, we compared COMB and CIT predictions with existing global trait maps generated through model-based extrapolations from sparse trait data, though using differing methodologies (Table 1). Three widely studied traits—specific leaf area (SLA), leaf N (mass), and leaf N (area)—were selected due to their broad overlap with prior trait mapping efforts.

At all resolutions and across all three traits, COMB predictions from this study exhibited stronger correspondence with independent, held-out vegetation survey data than previous trait maps. COMB models consistently produced the highest Pearson correlation coefficients (*r*), with values as high as 0.68 at 22 km resolution for leaf N (area). Even at coarser scales, COMB models retained high predictive accuracy, with average *r* values exceeding those of earlier studies. Notably, citizen science-only models (CIT), which employed no vegetation survey data during training, also demonstrated competitive performance, often outperforming previously published trait maps and achieving the second-highest correlations for most trait-resolution combinations.

### 2.6 Importance of environmental predictors

We assessed the relative importance of each Earth observation predictor to determine which predictor datasets contributed most to model performance (Fig. A.5). The predictor datasets included WorldClim bioclimatic variables (n=6), MODIS land surface reflectance (n=72), SoilGrids soil properties (n=61), VODCA vegetation optical depth (n=9), and global canopy height data (n=2) [19–22, 37].

For models trained with the combined dataset (COMB), MODIS surface reflectance emerged as the most influential predictor set overall (mean importance = 0.083), followed by WorldClim bioclimatic variables (0.051), SoilGrids soil properties (0.042), canopy height (0.008), and vegetation optical depth (0.006). Although MODIS, WorldClim, and SoilGrids contributed more strongly to trait predictions on average, the importance of individual predictors varied widely across traits. Notably, the highest single predictor importance was observed for a WorldClim variable (0.139, stem conduit density), followed by MODIS (0.091, specific leaf area), SoilGrids (0.025, rooting depth), canopy height (0.023, plant height), and vegetation optical depth (0.013, leaf N [mass]).

## 3 Discussion

Integrating diverse data sources, including structured vegetation surveys and opportunistic citizen science observations, provides a means to expand spatial coverage and improve trait predictions at a global scale. While citizen science data inherently contains noise and bias due to unstructured and opportunistic sampling [38–41], its broad geographic distribution and taxonomic coverage complement the systematic yet spatially limited nature of expert vegetation surveys. Models trained on a combination of these data sources (COMB) achieved predictive power comparable to those trained solely on vegetation surveys (SCI; Fig. 5), while also benefiting from markedly improved spatial transferability (Fig. 4).

Predictive performance was similar between SCI and COMB models, likely due to the down-weighting of citizen science samples in the COMB subset aimed at ensuring that models prioritized the more structured and reliable patterns from vegetation surveys. However, despite this down-weighting, the inclusion of citizen science observations still led to a notable reduction in the coefficient of variation and an expanded area of applicability. This reduction in uncertainty indicates that, while vegetation surveys offer higher-quality trait data, citizen science observations can enhance spatial generalizability and predictive confidence, particularly in regions with limited survey coverage. These benefits were especially pronounced in remote and under-sampled regions, such as alpine and boreal zones, where sparse vegetation surveys limit trait mapping efforts [25, 29]. Given the sensitivity of these ecosystems to climate change, permafrost dynamics, treeline shifts, and associated greening and browning trends, improving the spatial robustness of trait models is increasingly important [42].

The benefits of integrating multiple data sources were most pronounced at finer spatial resolutions (e.g., 1 km), where environmental conditions are more homogeneous and local-scale species assemblages are better captured. At these scales, the broader species occurrence data improved spatial coverage, expanding the area of applicability and enabling models to generalize to a wider range of environments. However, at coarser resolutions, the distinction between SCI and COMB models diminished, as spatial aggregation reduced variance in the predictor and trait spaces.

Model accuracy, however, followed a slightly different pattern (Fig. 5d). Some traits—such as leaf area, seed mass, and stem density—exhibited higher predictive accuracy at coarser resolutions while others plateaued or declined beyond 55 km resolution. This trend suggests that certain traits may align more strongly with large-scale climatic gradients, while others are driven by localized environmental and ecological factors, as well as land use and local management. These findings emphasize the need for trait-specific assessments of spatial scaling effects to refine predictions across different functional trait dimensions.

Trait maps generated using the combined data approach (COMB) exhibited higher accuracy compared to previous global trait mapping efforts when validated against independent community-weighted mean (CWM) trait data (Table 1). This improvement is likely due to the complementary nature of the input data sources, allowing COMB to leverage the strengths of both vegetation surveys and citizen science observations while mitigating their individual limitations. Notably, models trained exclusively on citizen science data (CIT), which never incorporated vegetation survey data, also outperformed most previously published trait maps. These findings support those of Wolf *et al.* [28], reinforcing the value of crowdsourced species observations in trait mapping. Despite the biases inherent in opportunistic citizen science data, the sheer volume and geographic coverage of these datasets appear to encode meaningful ecological patterns.

These improvements also likely reflect differing trait representation and underlying methodological assumptions and goals. For example, several previous trait mapping efforts have relied primarily on remote sensing proxies or broad-scale climate-trait correlations, which effectively capture dominant vegetation patterns but may not fully represent the trait variation present across plant communities. A key distinction of our approach is its ability to aggregate species-level trait data from multiple sources, allowing for a direct estimation of community trait composition rather than relying on aggregated plant functional type (PFT) classifications. Trait products that integrate discrete land cover or PFT information—particularly those derived from top-of-canopy remote sensing—have been shown to better reflect the traits of the most dominant functional groups within a given region rather than CWM traits [43]. Here, it was our aim to capture traits across the full vertical structure of plant communities, a perspective particularly relevant for applications in ecosystem process modeling, functional diversity assessments, and dynamic vegetation models [1, 12, 16].

To model each trait, 150 Earth observation and environmental predictors representing approx-imately 19 billion observations were utilized, prompting critical consideration of whether such a highly data-intensive approach is justified. Given the scale of these inputs, we calculated predictor importance using feature permutation to quantify the relative influence of a given predictor or predictor set on prediction quality (Fig. A.5).

MODIS surface reflectance was, on average, the most influential predictor set overall, followed by bioclimatic variables and soil properties, while canopy height and vegetation optical depth played a smaller role. These results align with prior research demonstrating that many functional traits are primarily structured along climate and, to a lesser extent, soil gradients, while their fine-scale variability may be better captured through optical remote sensing [1, 2, 44, 45]. Of course, such trends depend heavily on scale and the trait in question, and individual trait models exhibited distinct responses to different predictor types (Fig. A.6). Stem conduit density, for example, was most strongly influenced by a WorldClim bioclimatic variable, specific leaf area by MODIS reflectance, and rooting depth by SoilGrids. Further, canopy height and vegetation optical depth unsurprisingly contributed most to modeling structural traits like plant height and leaf width. These results suggest that no single environmental predictor is universally superior at predicting all plant functional traits and affirm that varied predictor data may be necessary to begin to encode the complexity of the biosphere and its functional diversity.

Despite the advantages of integrating multiple crowdsourced datasets, sampling biases remain a challenge. Vegetation surveys, citizen science data, and trait databases each exhibit geographic clustering and taxonomic gaps due to accessibility constraints, observer behavior, and historical research focus in the global north [38, 39, 46]. In African savannas, for example, grasses—despite being dominant across vast regions—are underrepresented in citizen science observations due to the difficulty of taking meaningful photographs of them, specialized identification requirements, and morphological similarities that complicate species-level classification. Likewise, vegetation surveys often target specific communities of interest rather than providing representative coverage at broader spatial scales. Addressing these biases will require targeted efforts to improve taxonomic and seasonal coverage in trait-mapping initiatives, and the rapid expansion of citizen science projects and increased collaboration among scientists present valuable opportunities to close these gaps in the near future [28, 29].

The influence of sampling density on predicted trait values is particularly evident in regions with sharp differences in observation coverage. In Portugal, for example, where CIT observation density is high but SCI surveys are sparse, trait patterns differ noticeably from those in neighboring Spain, where fewer CIT observations are available. This discrepancy likely stems from the inclusion of multiple country-wide forest inventories from Portugal in the GBIF database which may introduce a taxonomic bias in favor of woody plants. The resulting contrast in sampling density and methodology across the Iberian Peninsula’s unique environmental conditions may, in turn, confound model prediction, as indicated by the high coefficients of variation observed within Portugal compared to the relatively low variance in Spain and western France. These findings highlight the need for systematic bias corrections [41], such as spatial filtering or weighting schemes, to reduce observation-driven artifacts in trait predictions while also emphasizing the benefits of more comprehensive vegetation surveys.

Expanding citizen science and vegetation survey datasets enhances our understanding of plant community composition, but trait mapping remains constrained by the limited availability of in situ trait measurements. Most species observed in this study lacked corresponding trait records in the TRY database, with only 28% of citizen science observations and 43% of vegetation survey species records successfully matched (though ∼85% of all observations still fell within these species). This “Hutchinsonian shortfall” is by no means limited to this study, however, and despite ongoing database expansion and advanced gap-filling techniques, additional strategies are needed to bridge these gaps [25, 47, 48]. Computer vision, for instance, presents a promising opportunity to leverage the vast number of citizen science photographs to infer certain missing traits [49], not only improving trait coverage but also complementing traditional trait measurement efforts.

Even when species observations are successfully matched to trait databases, the “naive” approach of assigning species-mean trait values overlooks intraspecific trait variation (ITV) across environmental gradients. While ITV may contribute less to trait patterns at large spatial scales, it remains a key factor in functional diversity assessments [24, 50–52]. Importantly, the relevance of ITV varies by trait; for example, plant height exhibits substantial within-species variability, whereas specific leaf area (SLA) tends to be more stable [53]. Approaches such as incorporating in situ trait measurements or local PFT conditions within a given spatial radius may help capture ITV, though they often limit the number of species included per grid cell [45, 54]. Future trait-mapping efforts should explore integrating ITV-aware modeling approaches to improve ecological realism in global trait predictions.

The integration of diverse data sources in trait mapping provides a powerful framework for capturing plant functional diversity, yet several avenues remain for advancing these methodologies. One major advantage of our approach is its adaptability; beyond linking species observations to mean trait values, it allows for the estimation of trait distributions, including quantiles, ranges, and variance. This flexibility could also be leveraged to better represent the variability within specific plant functional types (PFTs), particularly for groups such as grasses and shrubs, where within-PFT variation is often masked by broad community-level averaging [43, 55]. Additionally, multi-label inference, an approach that enables models to predict multiple correlated traits simultaneously, could enhance predictive accuracy by leveraging known relationships among traits [56–59]. More extensive refinement of trait-specific predictor selection could also improve model efficiency by emphasizing the most relevant environmental variables for each trait. Trait predictions might be further improved by constraining species observations to those characteristic of specific vegetation formations, or PFTs, to ensure that trait-environment relationships are inferred within ecologically coherent units [2]. Beyond static trait predictions, future research could also focus on modeling key functional diversity (FD) and biodiversity metrics instead of solely community-weighted mean (CWM) traits. Understanding the global distribution of FD at high resolution may shed more light on the debated role of FD in ecosystem resilience and resistance to environmental change [30, 60–62].

Future advancements in remote sensing technologies offer significant opportunities for refining trait models. The increasing availability and planned expansion of hyperspectral imaging, satellite-derived canopy fluorescence, and high-resolution structural metrics will provide new avenues for detecting fine-scale trait variability [63–65]. Moreover, incorporating temporally resolved environmental predictors—such as ECMWF Reanalysis in combination with the high temporal resolution of MODIS and other remote sensing missions—could enable the development of multi-temporal trait maps, providing dynamic snapshots of how trait distributions respond to environmental fluctuations [66, 67]. By leveraging these time-series data, future trait models could better capture phenological shifts, disturbance responses, and other temporal dynamics that influence plant functional traits across different ecosystems. However, while environmental predictors may include high temporal resolution, the more restricted temporal character of trait measurements will likely remain a limiting factor.

## 4 Conclusion

This study demonstrates the potential of integrating multiple crowdsourced data repositories with high resolution Earth observation data to enhance global plant trait mapping. By combining expert vegetation surveys, citizen science-driven biodiversity observations, and trait measurement databases, our approach maximizes spatial coverage, trait representation, and predictive accuracy while mitigating individual data source limitations. However, these advancements were only made possible through the collective efforts of the scientific community and the broader public, facilitated by the growing framework of open, accessible, and reusable data. This collaborative, data-driven paradigm represents a fundamental shift in how we approach long-standing challenges in Earth system science. As curated expert data collections as well as crowdsourced biodiversity and trait databases continue to expand and data-sharing initiatives grow, future work should focus on improving bias correction, capturing trait variability, and integrating temporal dynamics to further refine functional trait predictions. By fostering continued collaboration between researchers, institutions, and citizen scientists, we can move toward a more comprehensive and globally representative understanding of plant functional diversity.

## 5 Methods

### 5.1 Trait data from the TRY Plant Trait Database

Data for 31 plant functional traits linked to key ecological processes (Table 2) and measured across 74,245 species were retrieved from the gap-filled TRY Plant Trait Database, version 5 (www.try-db.org) [14, 25, 68–71]. The TRY database is an aggregation of trait measurements from individual plants, often with multiple trait measurements per species. Despite providing robust mean trait values, the collection remains sparse and heterogeneous across environments and plant life stages in some areas, and so gap-filling utilizing Bayesian hierarchical probabilistic matrix factorization was implemented to expand its coverage [48, 68]. To minimize the impact of extreme values, outlier filtering was applied to each trait by excluding values below the 5th percentile and above the 95th percentile. Given the wide variation in functional trait distributions, trait data were transformed using Yeo-Johnson power transformation to standardize the distributions prior to model training [72].

### 5.2 Citizen science plant observations

Vascular plant species observations were retrieved from the Global Biodiversity Information Facility (GBIF; www.gbif.org) on 10 April 2024. Initial download parameters selected only observations that were members of Tracheophyta, included a geospatial reference, were marked as “present”, and were non-cultivated. Prior to filtering, the GBIF data included 339,971,350 observations of 314,217 plant species from 12,645 datasets across 113 countries [73]. Included in the GBIF data were nearly 100 million observations from popular citizen science initiatives, including more than 31 million observations from the iNaturalist Research-grade Observations dataset and over 24 million observations from Observation.org and Pl@ntNet combined [32, 33, 74]. Importantly, GBIF aggregates observations from more than just popular citizen science applications, but given their predominant presence in the database as well as the large variability among the other data sources, we chose to refer to the totality of GBIF as “citizen science” for ease of readability and accessibility. Before spatial aggregation (see details below), GBIF observations were randomly subsampled at each spatial resolution so that each resulting grid cell contained no fewer than 10 and no more than 500 observations.

### 5.3 Vegetation survey data

Vegetation survey data was retrieved from sPlot, a collation of worldwide vegetation plots including plot-level species relative coverage [29]. sPlot version 4.0 includes 52,942,365 occurrences of 91,603 species across 2,534,183 plots in 147 countries. Though GBIF observations cover a much larger geographic extent (Fig. 2), sPlot surveys have the advantage of incorporating an often more strategic sampling design, providing true absence information, and documenting species relative cover. Therefore, after matching with traits from TRY, sPlot can provide community-level trait statistics, such as community-weighted mean, median, and percentiles. This positions sPlot as a preferred alternative to GBIF in areas of overlapping or missing spatial coverage, as well as a benchmark against which trait extrapolation products can be compared.

### 5.4 Assigning trait values to citizen science observations and vegetation surveys

Following the methodology described by Wolf *et al.* [28], we calculated species-level means for the selected traits and assigned them to all matching species observations in both GBIF and sPlot using species name as the key matching criteria. Due to inconsistencies in species name formatting, all species identifiers in each dataset were first truncated to the first two words containing their primary binomial names and matched in a case-insensitive manner. Not all species present in GBIF and sPlot were present in TRY, and so some species were excluded (see Results), though similar matching efforts have shown that the species that remained still likely represent plant communities [29, 75]. For sPlot observations, plot-level community-weighted mean trait values were calculated by multiplying the relative cover of each species by the mean trait from TRY and averaging the resulting weighted values for each plot.

### 5.5 Spatial aggregation of citizen science and vegetation survey traits

In order to match the derived GBIF and sPlot trait values with gridded Earth observation data, we spatially aggregated both trait subsets. GBIF observation locations and sPlot plot locations were first projected from geographic coordinates into Equal-Area Scalable Earth (EASE) Grids, version 2.0, Global Cylindrical (EPSG: 6933). The GBIF individual trait values and sPlot plot-level community-weighted mean (CWM) trait values were each aggregated into separate 1 km × 1 km raster grids using the mean. As a result, the GBIF grids represented frequency-weighted mean trait values, which have been shown to approximate CWM traits [28], while the sPlot grids contained mean CWM values. For the combined trait data subset (COMB), GBIF and sPlot aggregations were merged, with sPlot values taking precedence in grid cells where both trait means from both datasets were present.

### 5.6 Gridded Earth observation data

#### 5.6.1 Bioclimatic variables

Climate data has been shown to largely drive the global variation in plant traits, and previous studies have successfully predicted plant traits via the environment [45, 54, 76–79]. More specifically, temperature influences photosynthesis by integrating energy availability from solar radiation. Plant productivity, reliant on photosynthesis, is also tied to water availability, making precipitation dynamics a crucial indicator of plant functional configuration [80]. Bioclimatic variables were extracted from the WorldClim database with a resolution of 0° 10^′^0^′′^ [22]. The following bioclimatic variables were selected: Annual Mean Temperature (BIO1), Temperature Seasonality (BIO4), Temperature Annual Range (BIO7), Annual Precipitation (BIO12), Precipitation of Wettest Month (BIO13), Precipitation of Driest Month (BIO14), and Precipitation Seasonality (BIO15). To calculate Precipitation Annual Range (BIO13-14) we subtracted Precipitation of Driest Month (BIO14) from Precipitation of Wettest Month (BIO13) and later discarded them, using only BIO13-14 to explain precipitation variability. As mentioned above, BIO1 and BIO12 are known predictors of plant traits. Likewise, climate variables associated with range and seasonality have also been shown to correlate with plant traits. Therefore, we specifically chose the additional four climate variables (BIO4, BIO7, BIO13-14, BIO15) to capture the annual variations in the site-specific climatic conditions.

#### 5.6.2 Surface reflectance

Given that plant canopies are configured to interact with light, their optical reflectance properties can inform their functional trait configuration [45, 81]. When considering the prospect of perpetually-updated plant trait models, remote sensing data is especially appealing given its high spatial and temporal availability. For this study, surface reflectances in the visible and infrared spectra were collected from the MODIS MOD09GA v061 (MODIS/Terra Surface Reflectance Daily L2G Global 1 km and 500 m SIN Grid) dataset [19]. MODIS was selected due to its more reliable and continuous large-scale coverage when compared to other optical remote sensing missions like Landsat and Sentinel-2. Bands 1-5 (620-670 nm, 841-876 nm, 459-479 nm, 545-565 nm, and 1230-1250 nm, respectively) were used, as well as an NDVI band which was calculated using bands 1 (red) and 2 (near-infrared). In order to preserve as much spatial and temporal information as possible, surface reflectances for each band were obtained at 1 km resolution and temporally aggregated into 12 monthly averages spanning 20 years from March 2000 through March 2020, yielding 72 total predictors (12 × 5). These operations were performed using Google Earth Engine.

#### 5.6.3 Soil properties

Because soil properties influence nutrient and water availability—key factors in plant resource acquisition and trait composition—we included high-resolution soil data under the assumption that it could help explain some of the trait variance [48]. Mean soil properties were obtained from the SoilGrids 2.0 dataset—global soil property predictions at six standard depth intervals at a spatial resolution of 250 meters—produced and hosted by the International Soil Reference and Information Centre (ISRIC) [21]. All available soil properties at all depth intervals were used, including pH, soil organic carbon content, bulk density, coarse fragments content, sand content, silt content, clay content, cation exchange capacity, total nitrogen, soil organic carbon density, and soil organic carbon stock.

It is important to note that the predictor datasets used during model training for SoilGrids predictions include MODIS optical observations (middle and near-infrared bands) as well as WorldClim 2.0 bioclimatic variables, and therefore some collinearity may exist between the predictors used in this study.

#### 5.6.4 Vegetation optical depth

Vegetation optical depth (VOD) measures the opacity of vegetation to microwave radiation, indicating the amount and water content of plant biomass across landscapes. In addition to plant water content, VOD can inform ecosystem properties relevant to plant traits such as vegetation density, biomass, and environmental conditions [82, 83]. Further, VOD can discriminate between vegetation types, a reliable predictor of plant traits [2, 27, 45, 54, 84]. Global, multi-sensor measurements of VOD have been made available as part of the global long-term microwave Vegetation Optical Depth Climate Archive (VODCA) [20]. We obtained all available measurements in the C, Ku, and X bands from all available years (C-band: 2002-2018; Ku-band: 1987-2017; X-band: 1997-2018) at a native spatial resolution of 0.25° and upsampled all measurements to 1 km using bilinear interpolation. Bilinear interpolation was utilized over nearest-neighbor upsampling (NN) as NN tended to result in the imprinting of the 0.25° grid cells during model extrapolation. Three temporal aggregations were then computed across each band time series: mean and the 5th and 95th percentiles.

#### 5.6.5 Canopy height

Canopy height not only has an obvious direct relationship with community-level plant height, but has also been shown to be correlated with above-ground traits like stem diameter, stem specific density, and seed mass, as well as below-ground and other hydraulic traits such as stem conductivity and potential root length and mass [56, 85–87]. Additionally, taller plants directly affect the competitive landscape by reducing light availability to below-canopy communities as well as demanding more water and nutrients than smaller individuals due to increased transpiration and photosynthesis [88, 89]. To represent global canopy height, we utilized the global canopy height and canopy height standard deviation products developed by Lang *et al.* [37] due to their uniquely high spatial resolution compared to other canopy height models.

#### 5.6.6 Geospatial reprojection, resampling, and masking

All predictor datasets, with the exception of VOD, were obtained at resolutions similar to or finer than 1 km, and so each dataset was first reprojected to EASE (EPSG: 6933) for standardized, area-consistent global referencing and then spatially resampled using weighted mean resampling to 1 km (∼0.01°), 22 km (∼0.2°), 55 km (∼0.5°), 111 km (∼1°), and 222 km (∼2°) grids. As mentioned above, upsampling was required for VOD observations in order to be matched with other predictors and trait grids at 1km and 22 km resolutions. Lastly, water and urban land cover pixels were masked from all predictor maps using ESA WorldCover 10m v100 [90].

### 5.7 Model training and validation

#### 5.7.1 Matched predictors and trait data subsets

Prior to model training, environmental predictors and GBIF and sPlot trait data (labels) were matched by pixel coordinates and converted to tabular format. Though our machine-learning approach could tolerate missing values, pixels containing fewer than 60% of the predictors were excluded from the final training sets. Overall, the three subsets of trait data were matched with environmental predictors: vegetation survey trait community-weighted means (CWM) only (SCI), citizen science trait means only (CIT), and a combination of both, where vegetation survey CWMs were preferred when both were present (COMB; Table 3). This approach was repeated to generate training data for models at each resolution.

**Table 3:**
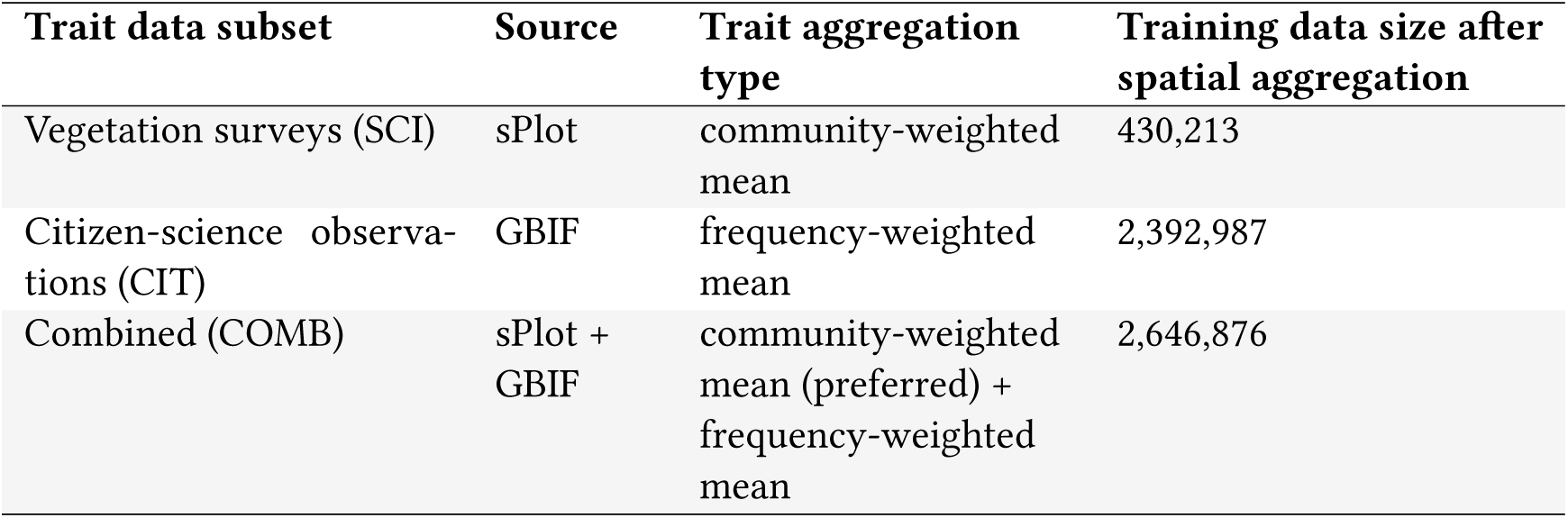
Trait data subsets used for model training. Models for all traits were trained for each trait data subset at multiple resolutions and evaluated against held-out, spatially independent community-weighted mean traits using spatial K-fold cross-validation. In the case of the COMB trait data subset, SCI and CIT were merged with a preference for SCI values when both SCI and CIT data were present.

#### 5.7.2 Model training and spatial cross-validation

After matching by pixel coordinates, models were trained for all traits across each trait data subset using LightGBM gradient-boosting decision trees [91]. LightGBM utilizes a suite of boosted decision trees and is capable of capturing complex, nonlinear relationships among high-dimensional tabular data while avoiding overfitting and remaining resource-efficient [35]. Additionally, Light-GBM is capable of reasonably handling missing values by ignoring them during splitting, then allocating them to whichever side reduced loss the most. This was necessary as not all predictors had complete spatial overlap, and so, while observations that had more than 40% missing predictor values were filtered prior to training, some missing predictor values persisted depending on the spatial coverage of each particular trait.

When validating machine learning-based geospatial models, spatial autocorrelation must be considered [92, 93]. Traditional validation methods such as random K-fold or leave-one-out cross-validation (CV) tend to produce overly optimistic results because they do not account for spatial clustering in training data or discrepancies between training and extrapolation feature spaces [92, 94]. While probability sampling and design-based inference may be preferable for reference datasets with sufficiently broad spatial coverage, spatial K-fold cross-validation (SKCV) has been shown to be a robust alternative for sparse, spatially clustered datasets [95]. When combined with supplementary quality indicators—such as the coefficient of variation and area of applicability (AOA), discussed below—SKCV provides a more reliable assessment of model generalizability [36, 92, 96].

To construct SKCV folds, we first determined the spatial autocorrelation range of sPlot community-weighted mean (CWM) trait values by computing spherical semivariograms at a 1 km resolution. We then overlaid a hexagonal grid with cells sized according to this range and randomly assigned each cell to one of five folds. All trait observations from both sPlot and GBIF were assigned a fold ID based on their respective hexagons. To ensure balanced fold distributions, we ran 100 simulations of random fold assignments and selected the final configuration based on minimizing dissimilarity between folds. Dissimilarity was quantified using Kolmogorov-Smirnov tests, with the chosen fold assignment maximizing the mean *p*-value, thereby minimizing the likelihood of systematic differences between folds.

Model training and validation was performed using the leave-one-out (LOO) method, in which predictor-trait pairs from a single fold were set aside for model validation while fitting was performed on the remaining data, and the process was repeated for each remaining fold. All trait model variants (SCI, CIT, and COMB), were validated using vegetation survey community-weighted mean trait values from the held-out fold to ensure validation integrity and spatial independence of the validation data. Due to the significant imbalance in sample size between GBIF and sPlot pixels after spatial aggregation, samples in the combined trait data subset (COMB) were weighted based on their origin. sPlot vegetation surveys, which represent community-weighted means, were deemed more reliable than citizen science observations. Consequently, weights were assigned to GBIF observations inversely to their proportion relative to sPlot observations in the combined training set. Additionally, feature pruning was performed during model training based on automated sensitivity analysis via permutation feature importance. Features were dynamically removed during model ensemble creation if their removal improved model performance. Because final “models” were actually an ensemble of child models, not all child models necessarily utilized the same pruned feature set. Finally, models used to produce final trait prediction maps were trained on all available environmental predictor data.

#### 5.7.3 Predictor importance

The retrieval of predictor importance, or the weight given to a particular feature during model training, is an important aspect of model interpretability. Feature and dataset importance were calculated using the permutation importance method, in which the values of each feature or set of features (dataset) are randomly shuffled and model performance is measured [97]. The permutation importance is quantified as the performance difference when prediction is performed on data containing the shuffled feature. Because we implemented SKCV, feature importance was defined as the average feature importance across all CV folds.

### 5.8 Model performance and spatial transferability

Citizen science trait values, given their opportunistic provenance, are noisy and biased [39]. As mentioned above, we validated all model runs against spatially independent sPlot community-weighted mean trait values that were not used during model training for each of the three trait data subsets (CIT, SCI, and COMB). Model performance was assessed using average normalized root mean squared error (*nRMSE*) and Pearson’s correlation coefficient *r* based on the combined observed versus predicted values of all spatial cross-validation runs. To further assess model robustness, we calculated the pixel-wise coefficient of variation (COV) by generating predictions from each cross-validation model, stacking the results, and computing the COV for each pixel along the vertical stack as the ratio of the standard deviation to the mean prediction. All above statistics were also assessed across biomes. Biomes were first retrieved from the Terrestrial Ecoregions of the World map and then consolidated from 14 biomes into 7 biome categories [98]. The biome-to-biome-category mapping can be found in the appendix (Table A.2).

#### 5.8.1 Area of applicability

While SKCV is useful for minimizing spatial dependence on model error estimation, it may not be sufficient for explaining the transferability of model extrapolations in areas where predictor data vary significantly from the reference data [40, 94, 99]. In particular, we were interested in assessing whether the abundance and coverage of citizen science observations impart greater spatial transferability to citizen-science based models (COMB) compared to using less-abundant vegetation surveys alone. To address this, we applied the methods developed by Meyer & Pebesma [36] for determining the dissimilarity index (DI) between predictor data used during model training (“train”) and unseen predictor data used for extrapolation (“new”), and thus the area of applicability (AOA) of the final trait maps. Briefly, dissimilarity in the predictor space was calculated by first computing the average minimum distance between observations in each cross-validation fold in “train” (i.e. the minimum distances between points in one fold from points in all other folds), followed by calculating the minimum distances between observations in the “new” data from the training data. The DI was then determined as “new” distances divided by mean “train” distance. In line with existing literature, we defined the DI threshold as of the cross-validated training data, with values below the threshold being within the AOA and values above the threshold (outliers) being outside it. DI and AOA were computed for all models at all resolutions [36]. To quantify whether the inclusion of GBIF data increased the AOA of trait models, we compared the AOA of all models across all training sets and calculated the change in overall extent.

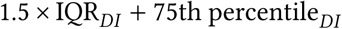

It should be noted that the methods described by Meyer & Pebesma [36] do not address the handling of missing values. Because LightGBM can tolerate missing values and we did not do any data imputation prior to training, we had several options when calculating AOA: a) drop all observations in “new” and “train” that contained any missing features (and therefore misrepresent the actual spatial coverage of the training data as well as reduce feature space variance present in DI calculation compared to actual reference data used in training), likely resulting in an overly pessimistic AOA; b) drop features in both the “new” and “train” data that contained any missing values, likely resulting in an overly optimistic AOA as feature-space complexity is reduced; or c) a “middle ground” approach of imputing missing values and assuming that the resulting predictor space is still a robust representation of the true reference data. To retain as much of the original signature of the true reference data when calculating dissimilarity, as well as to ensure that final AOA maps matched the geographic extent of the predictions, we chose option “c”. However, we recognize that, given the novelty of this method, there is likely room for improvement. Missing value imputation was performed using the `NaNImputer` method from the Python “verstack” library, which fits LightGBM regression models for each feature to fill missing values [100].

#### 5.8.2 Trait map creation

We generated continuous trait maps for all traits at all resolutions for the SCI and COMB trait subsets as stacked rasters containing the inference data, COV, and AOA mask. The rasters were written as cloud-optimized GeoTIFFs in EPSG:6933 and the inference and COV bands were quantized to signed 16-bit integer format.

#### 5.8.3 Comparison of existing products against sPlot community-weighted means

In order to compare our trait maps with existing trait map products, we chose to use sPlot gridded CWMs as a benchmark. Trait maps from nine existing products covering three foliar traits—specific leaf area, leaf nitrogen per area (leaf N [area]), and leaf nitrogen per mass (leaf N [mass])—were selected and spatially aligned with sPlot CWMs at the nearest common resolution (1 km, 22 km, 55 km, 111 km, or 222 km). Most products were originally created at a single native resolution, but to enable performance comparisons across scales, they were downsampled to coarser resolutions when necessary. Upsampling to finer resolutions was avoided to prevent unnecessary data manipulation. The final trait values were transformed using a Yeo-Johnson power transform with the lambda values determined in the original transformation of the sPlot gridded CWMs. The Pearson’s correlation coefficient *r* between the transformed trait products and matched sPlot CWMs was then used to describe agreement. The specific foliar traits were selected due to their general commonality between the selected previous studies.

## 6 Data availability

The trait products can be obtained and visualized using the following resources:

- Data page: https://geosense-freiburg.github.io/global-traits/
- Online map viewer: https://global-traits.projects.earthengine.app/view/global-traits
- Download link: TBA

## 7 Code availability

- GitHub: TBA

## 8 Acknowledgments

This study was funded by the German Research Foundation (DFG) within the framework of Big-PlantSens (Assessing the Synergies of Big Data and Deep Learning for the Remote Sensing of Plant Species; project no. 444524904) and PANOPS (Revealing Earth’s plant functional diversity with citizen science; project no. 504978936). The study is supported by the TRY initiative on plant traits (http://www.try-db.org) and the sPlot consortium (http://www.idiv.de/splot). The TRY initiative and database are hosted, developed, and maintained by J.K. and G. Boenisch (Max Planck Institute for Biogeochemistry, Jena, Germany), currently supported by Future Earth/bioDISCOVERY and the German Centre for Integrative Biodiversity Research Halle-Jena-Leipzig (iDiv, DFG-FZT 118, 202548816). The sPlot is a strategic project of iDiv and is supported by the German Research Foundation (DFG-FZT 118, 202548816). F.M.S. gratefully acknowledges the support of the Italian Ministry of University and Research, under the Maria Levi Montalcini programme. S.W. was funded by the German National Research Data Infrastructure for Biodiversity, NFDI4Biodiversity, a DFG project, project no. 442032008. F.G.’s surveying efforts were funded by the Swiss National Science Foundation Postdoctoral Fellowships (TMPFP2 217531). F.R. acknowledges the support of Coordenação de Aperfeiçoamento de Pessoal de Nível Superior (CAPES) for postdoctoral fellowships (N°88887.974741/2024-00). This work would not have been possible without the contributions of ecologists and vegetation surveyors who diligently sampled field plots, curated datasets, and shared them through accessible databases. We also acknowledge the vital role of citizen scientists engaged in platforms such as iNaturalist and Pl@ntNet, whose time, observations, and local knowledge have been crucial in assembling high-quality, research-ready data.

## A Appendix

### A.1 Model performance

**Figure A.1:**
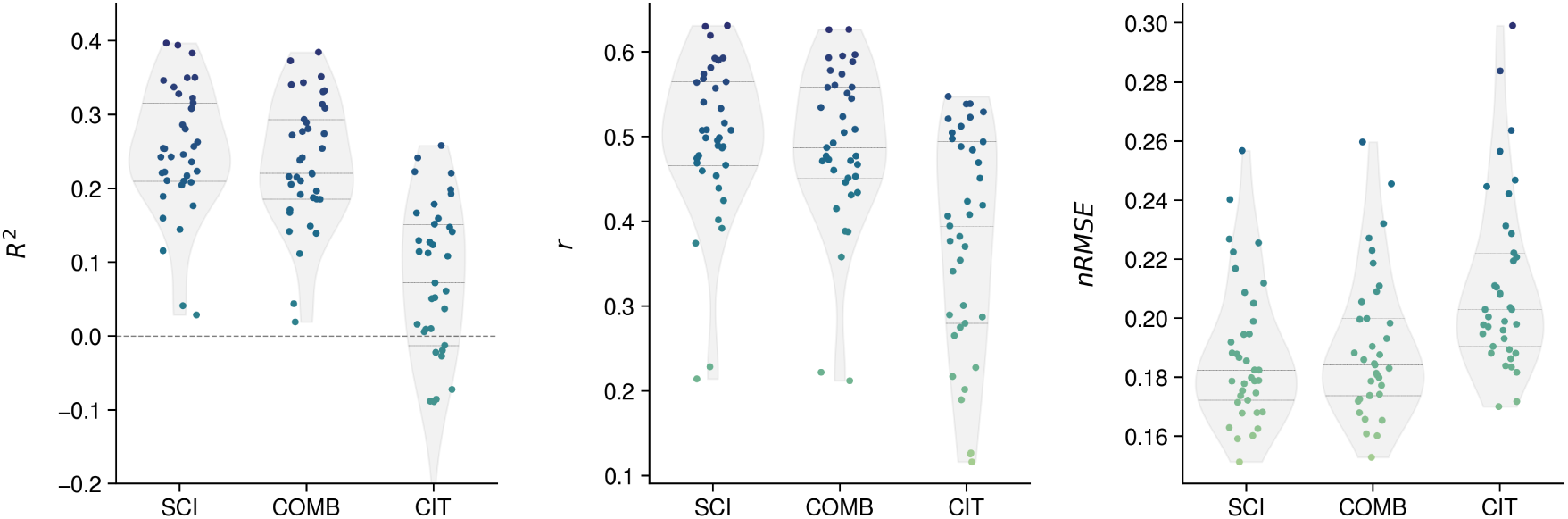
*R*^2^, Pearson correlation coefficient *r*, and normalized root mean squared error *nRMSE* of all traits for each trait subset: vegetation surveys (SCI), citizen science observations (CIT), and both sources combined (COMB). Metrics are the result of validation against spatially independent, unseen vegetation survey community-weighted mean trait values. The points indicate individual trait model performance, while the encompassing violin plots represent the performance distribution for each trait subset. Inner horizontal bars represent the 75th, 50th (median), and 25th quantiles. For *R*^2^, the horizontal line at 0 indicates the cutoff, wherein models below the line had a negative *R*^2^ (only present for some traits based on CIT).

### A.2 Spatial transferability

### A.3 Feature importance

### A.4 Biome definitions

**Figure A.2:**
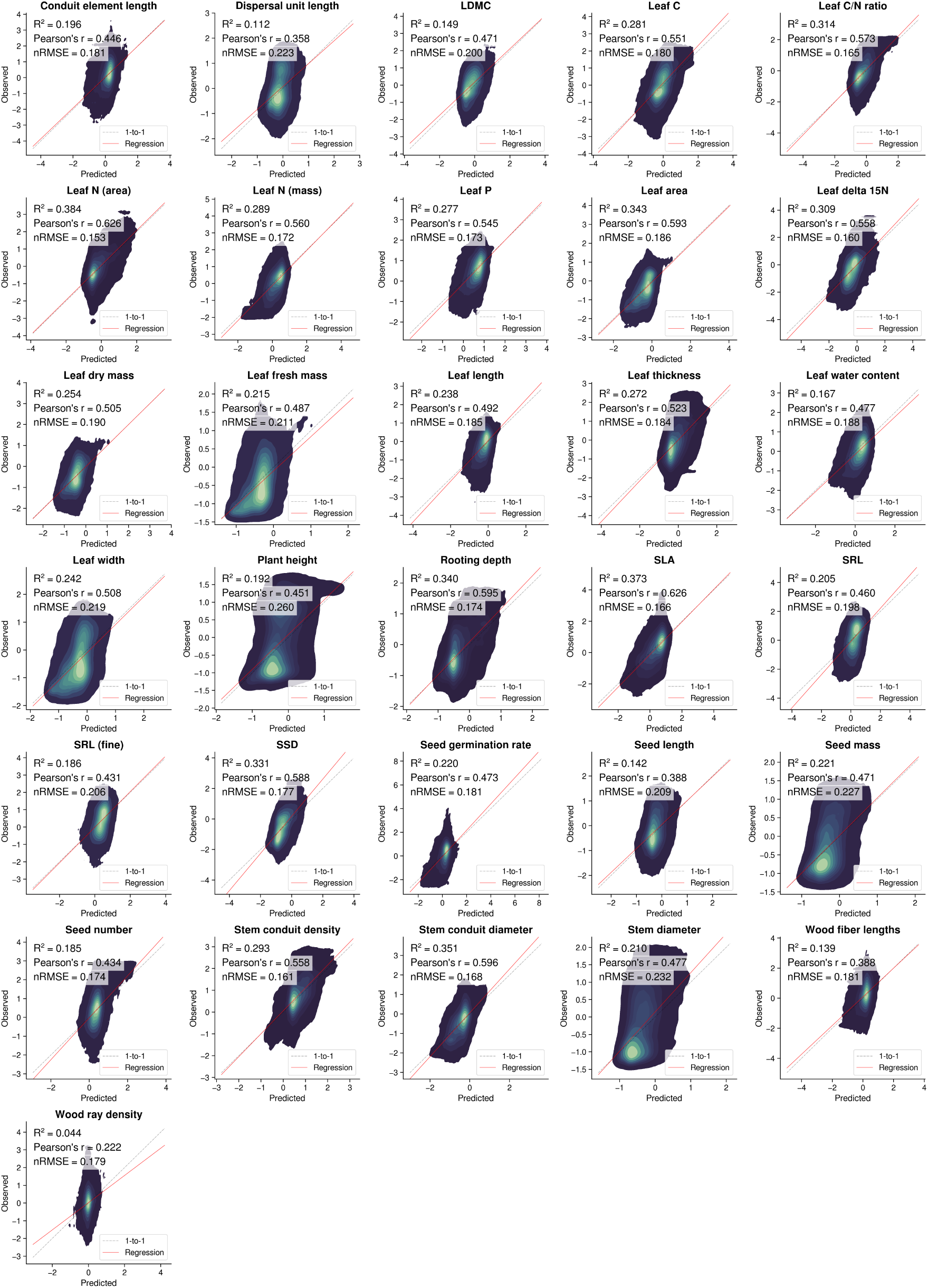
Observed versus predicted community-weighted mean trait values from models trained on vegetation survey and citizen science data (COMB). Scatterplots are presented as density plots, with lighter colors representing higher point density. Model performance statistics are shown, as well as a grey dashed 1-to-1 line representating a correlation of 1 and a linear regression line in solid red. All trait values are in Yeo-Johnson power-transformed space.

**Figure A.3:**
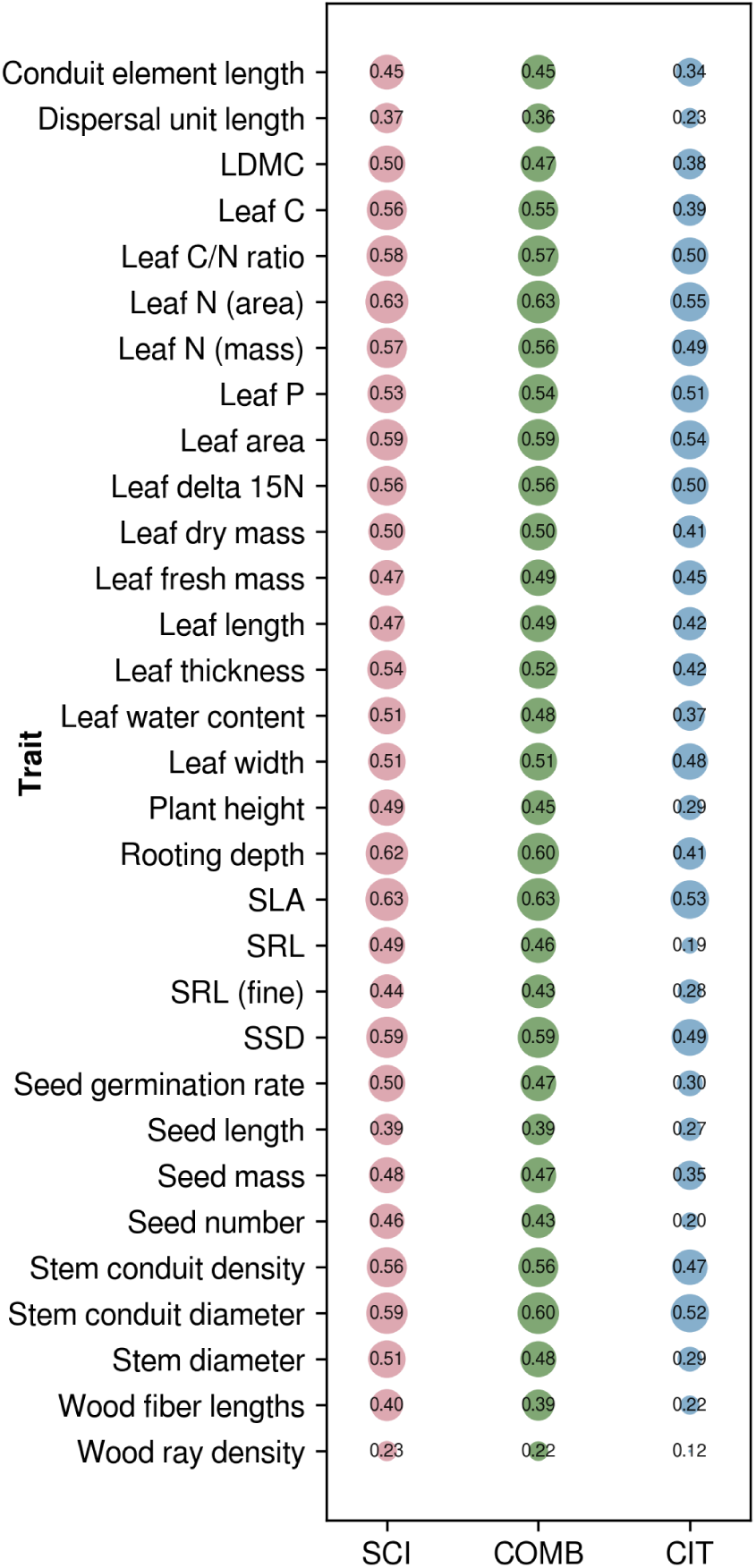
Pearson’s correlation coefficient *r* for all traits across each trait data subset at 1 km resolution. Circle size corresponds to the magnitude of *r*, with a minimum value of 0.12 (citizen science data only [CIT], wood ray density) and a maximum of 0.63 (shared by vegetation survey data [SCI] and combined trait sources [COMB], leaf N (area) and SLA). Circle size was normalized to the data range for visual clarity of differences in correlation.

**Figure A.4:**
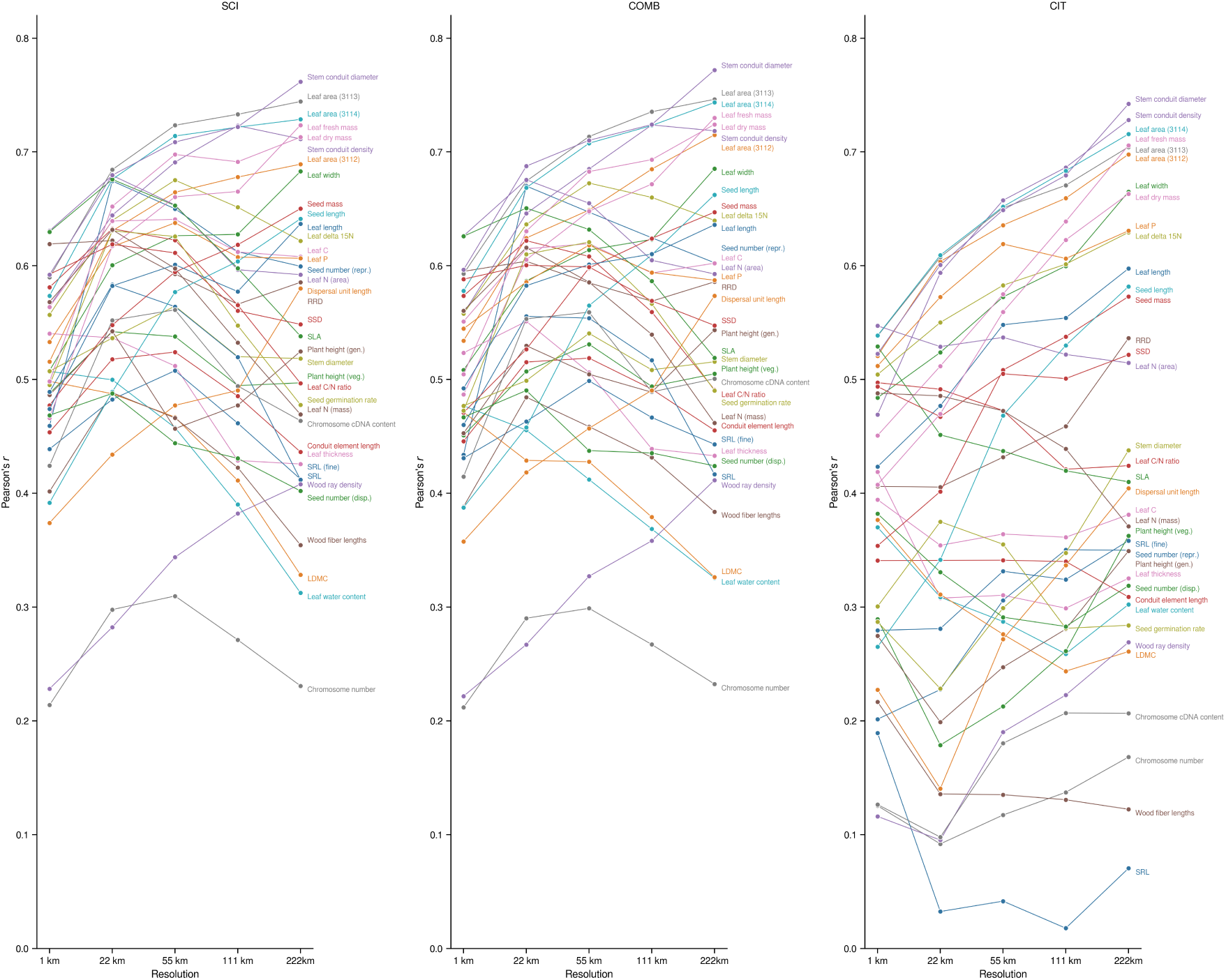
Pearson’s correlation coefficient *r* based on spatial K-fold cross-validation for SCI, COMB, and CIT models across all resolutions.

**Figure A.5:**
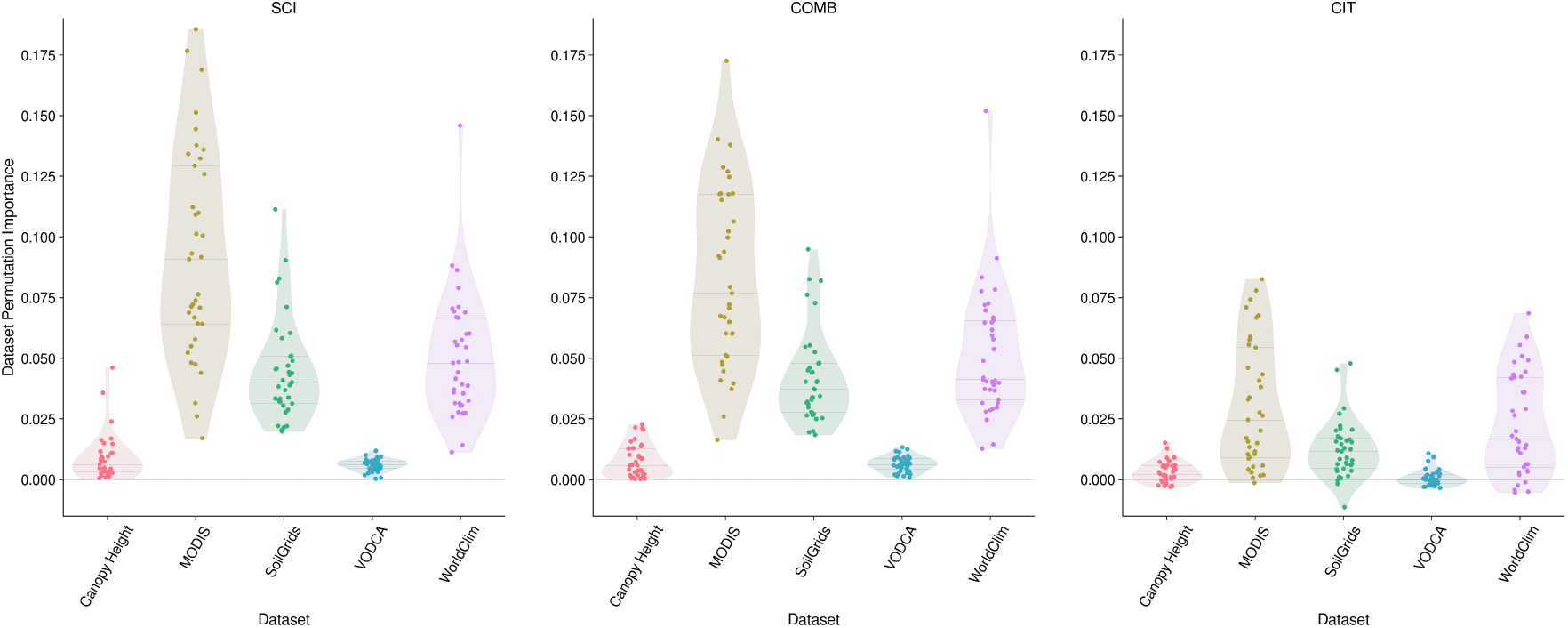
Dataset permutation importance of each dataset used in model training. Here, dataset permutation importance refers to the effect of randomly shuffling the values of *all* features within a given dataset on trait model performance. Individual dots represent traits, and each plot represents a different trait data subset: vegetation survey data alone (SCI, left), citizen science data alone (CIT, right), and a combination of both (COMB, middle).

**Table A.1:** Difference in area of applicability (AOA) between models trained on vegetation surveys supplemented with citizen science data (COMB) and models trained on vegetation survey data alone (SCI). Positive values indicate the extent to which COMB models resulted in a greater AOA. The largest increases for each resolution are highlighted in bold.

| Trait | 1 km | 22 km | 55 km | 111 km | 222 km |
| --- | --- | --- | --- | --- | --- |
| Chromosome cDNA content | 1.79% | 1.07% | 1.10% | 1.23% | 0.78% |
| Chromosome number | 2.27% | 1.86% | 1.38% | 1.20% | 0.93% |
| Conduit element length | 7.34% | 1.26% | 1.39% | 0.47% | 0.62% |
| Dispersal unit length | 0.41% | 0.53% | 0.44% | 0.49% | 0.35% |
| LDMC | 3.77% | 1.47% | 2.00% | 1.24% | 1.97% |
| Leaf C | 2.58% | 1.19% | 0.99% | 1.21% | 0.50% |
| Leaf C/N ratio | 4.89% | 2.17% | 1.79% | 1.26% | 1.85% |
| Leaf N (area) | 1.69% | 0.64% | 0.68% | 1.00% | 1.49% |
| Leaf N (mass) | 2.89% | 1.60% | 1.60% | 1.16% | 1.45% |
| Leaf P | 2.23% | 0.63% | 0.72% | 0.43% | 0.23% |
| Leaf area (3112) | 0.34% | 0.32% | 0.26% | 0.19% | 0.23% |
| Leaf area (3113) | 1.43% | 0.46% | 0.41% | 0.49% | 0.31% |
| Leaf area (3114) | 2.73% | 0.42% | 0.35% | 0.32% | 0.19% |
| Leaf delta 15N | 1.64% | 0.52% | 0.33% | 0.34% | 0.15% |
| Leaf dry mass | 3.42% | 0.61% | 0.40% | 0.38% | 0.27% |
| Leaf fresh mass | 0.47% | 0.06% | 0.12% | 0.15% | 0.15% |
| Leaf length | 0.49% | 0.57% | 0.21% | 0.13% | 0.27% |
| Leaf thickness | 1.78% | 2.49% | 0.55% | 0.26% | 0.39% |
| Leaf water content | 2.87% | 1.32% | 1.38% | 1.35% | 1.37% |
| Leaf width | 1.65% | 0.34% | 0.24% | 0.22% | 0.04% |
| Plant height (gen.) | 0.70% | 1.00% | 1.26% | 1.55% | 0.43% |
| Plant height (veg.) | 0.15% | 0.73% | 0.64% | 0.81% | 0.43% |
| RRD | 1.96% | 1.35% | 0.88% | 1.04% | 0.86% |
| SLA | 1.87% | 1.81% | **2.27%** | 1.10% | 1.85% |
| SRL | 0.82% | 1.89% | 0.83% | 1.81% | 1.93% |
| SRL (fine) | 7.62% | **3.21%** | 1.26% | 1.01% | 1.25% |
| SSD | 0.97% | 0.97% | 0.68% | 0.82% | 0.70% |
| Seed germination rate | 3.22% | 0.62% | 1.06% | 1.34% | **2.61%** |
| Seed length | 0.68% | 0.59% | 0.78% | 0.49% | 0.58% |
| Seed mass | 3.00% | 1.19% | 0.82% | 0.71% | 0.66% |
| Seed number (disp.) | 2.52% | 0.66% | 0.49% | 0.47% | 0.70% |
| Seed number (repr.) | 2.27% | 0.84% | 1.12% | **2.10%** | 0.43% |
| Stem conduit density | 1.19% | 1.12% | 0.50% | 0.57% | 0.31% |
| Stem conduit diameter | 2.08% | 0.85% | 1.00% | 0.52% | 0.50% |
| Stem diameter | 2.75% | 0.24% | 0.57% | 0.31% | 0.50% |
| Wood fiber lengths | **9.22%** | 2.66% | 1.07% | 0.84% | 0.78% |
| Wood ray density | 2.08% | 0.86% | 1.19% | 1.73% | 0.62% |

**Figure A.6:**
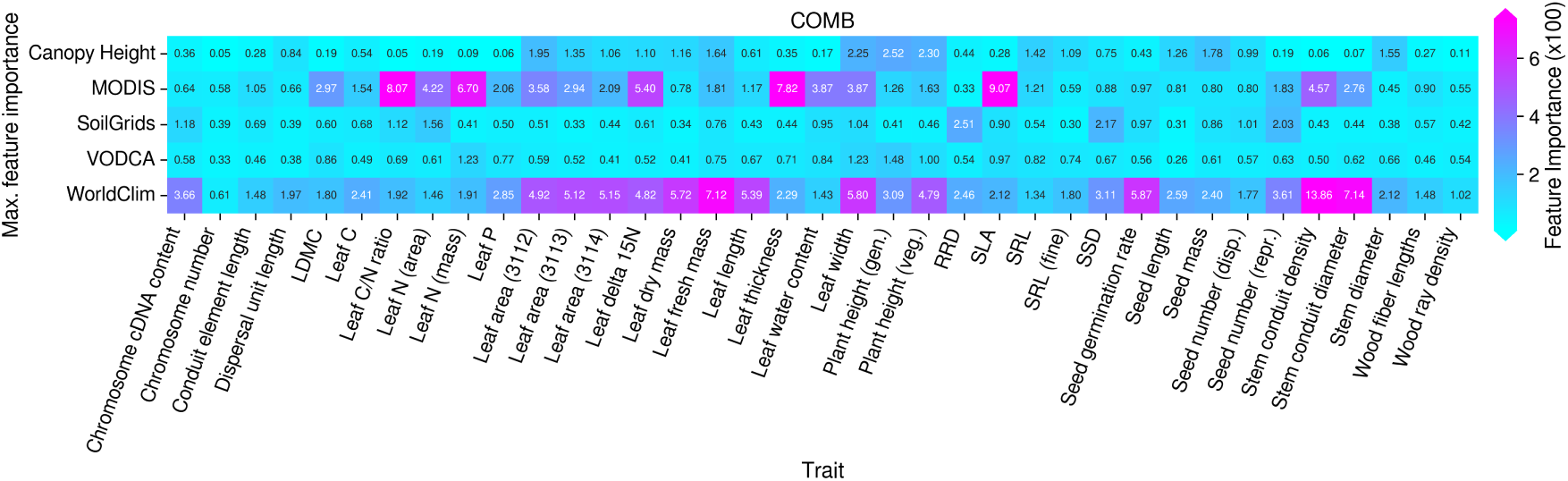
Maximum feature permutation importance of each dataset used in COMB model training. Here, maximum feature permutation importance refers to the maximum permutation importance of a feature within each dataset for each trait model. Values have been multiplied by 100 for readability.

**Table A.2:** Condensed biome definitions extracted from the Terrestrial Ecoregions of the World dataset by Olson *et al.* [98] and used for assessing model performance and spatial transferability across biomes.

| Biome Set | Original Biome from TEOW |
| --- | --- |
| Tropical/Subtropical Forests | Tropical & Subtropical Moist Broadleaf Forests |
|  | Tropical & Subtropical Dry Broadleaf Forests |
|  | Tropical & Subtropical Coniferous Forests |
| Temperate/Boreal Forests | Temperate Broadleaf & Mixed Forests |
|  | Temperate Conifer Forests |
|  | Boreal Forests/Taiga |
| Grasslands/Savannas | Tropical & Subtropical Grasslands, Savannas & Shrublands |
|  | Temperate Grasslands, Savannas & Shrublands |
| Mediterranean | Mediterranean Forests, Woodlands & Scrub |
| Deserts | Deserts and Xeric Shrublands |
| Wetlands | Flooded Grasslands and Savannas |
|  | Mangroves |
|  | Lake |
| Alpine/Polar | Montane Grasslands and Shrublands |
|  | Tundra |
|  | Rock and Ice |

